# Discovery of the gut microbial enzyme responsible for bilirubin reduction to urobilinogen

**DOI:** 10.1101/2023.02.07.527579

**Authors:** Brantley Hall, Sophia Levy, Keith Dufault-Thompson, Glory Minabou Ndjite, Ashley Weiss, Domenick Braccia, Conor Jenkins, Yiyan Yang, Gabi Arp, Stephenie Abeysinghe, Madison Jermain, Chih Hao Wu, Xiaofang Jiang

## Abstract

The degradation of heme and the interplay of its catabolic derivative, bilirubin, between humans and their gut microbiota is an essential facet of human health. However, the hypothesized bacterial enzyme that reduces bilirubin to urobilinogen, a key step that produces the excretable waste products of this pathway, has remained unidentified. In this study, we used a combination of biochemical analyses and comparative genomics to identify a novel enzyme, BilR, that can reduce bilirubin to urobilinogen. We delineated the BilR sequences from other members of the Old Yellow Enzyme family through the identification of key residues in the active site that are critical for bilirubin reduction and found that BilR is predominantly encoded by Firmicutes in the gut microbiome. Our analysis of human gut metagenomes showed that BilR is a common feature of a healthy adult human microbiome but has a decreased prevalence in neonates and IBD patients. This discovery sheds new light on the role of the gut microbiome in bilirubin metabolism and highlights the significance of the gut-liver axis in maintaining bilirubin homeostasis.

## Introduction

Bilirubin is an intermediate metabolite in the heme degradation pathway, a crucial facet of human physiology^1^. The homeostasis of molecules like bilirubin, cholesterol and bile acids is maintained by the gut-liver axis^2,3^. This interface between the gut, gut microbes, and liver, serves as one of the primary routes for nutrient absorption and waste secretion in the body. Similar to cholesterol and bile acids, bilirubin is secreted from the liver into the gut where the gut microbiota can metabolize it, affecting its reabsorption^4–7^. The homeostasis of serum bilirubin has significant impacts on human health with moderate levels of serum bilirubin having potential health benefits due to its antioxidant properties^8,9^, while high levels of serum bilirubin can be toxic leading to jaundice and in extreme cases kernicterus, a type of bilirubin-induced neurological damage^10^. Bilirubin metabolism is a vital component of heme metabolism and the gut-liver axis and it is essential to understand what microbes and enzymes are involved.

The gut microbiota is implicated in serum bilirubin homeostasis through the conversion of the bilirubin secreted into the gut to more soluble urobilinoids that can be readily excreted rather than entering enterohepatic circulation. While the importance of bilirubin enterohepatic circulation to serum bilirubin levels was suggested in the 1960s and 1970s^11,12^, it was largely undervalued until the work of Vitek et al. in the early 2000s, which showed the direct impact of microbial bilirubin metabolism on serum bilirubin levels in rats^13,14^. The gut microbiota has been shown to metabolize bilirubin in two ways: 1) the deconjugation of bilirubin-glucuronide molecules and 2) the reduction of unconjugated bilirubin to urobilinogen and subsequently stercobilinogen^8,14,15^. The deconjugation of bilirubin-glucuronide by human and bacterial beta-glucuronidases results in the production of unconjugated bilirubin which can be readily reabsorbed, leading to elevated serum bilirubin levels^16,17^. The reabsorption of unconjugated bilirubin can be limited by the microbial reduction of bilirubin to urobilinogen, which is excreted as a waste product in urine and feces, completing the heme excretion pathway^18,19^.

Bacteria are solely responsible for the reduction of bilirubin to urobilinogen, likely utilizing the molecule as a terminal electron acceptor in the anaerobic environment of the gut. Despite their recognized role in this process, the bacterial enzyme that reduces bilirubin to urobilinogen, hereafter called bilirubin reductase, has remained undiscovered^20^. Multiple bilirubin reducing bacteria have been identified including strains of *Clostridioides difficile^14^, Clostridium ramosum^21^, Clostridium perfringens^14^*, and *Bacteroides fragilis^8^*. Unfortunately, much of this work took place before genome sequencing was widely available and many of the strains are currently unavailable. Additionally, numerous changes to bacterial taxonomy cloud the true taxonomic identity of these species. Without knowledge of the gene encoding bilirubin reductase, it is difficult to draw conclusions about how gut microbial bilirubin metabolism affects serum bilirubin homeostasis, leaving a significant gap in our understanding of the heme excretion pathway^22–24^.

In this study, we aimed to identify the enzyme(s) responsible for bilirubin reduction to urobilinogen and perform a systematic evaluation of the distribution of these genes across bacterial taxa so that we could apply this knowledge to understand the relationship between bilirubin reduction and human health. We first screened multiple bacterial strains using a fluorescence assay and metabolomics, confirming that some known strains were able to reduce bilirubin. We also identified multiple new bilirubin reducing strains as well taxonomic relatives that do not reduce bilirubin. Combining comparative genomics between bilirubin reducing and non-reducing strains and biochemical inference of the putative bilirubin reductase reaction mechanism, we were able to identify a candidate bilirubin reductase gene we named *bilR*. We expressed *bilR* in *Escherichia coli* confirming its bilirubin reductase activity through fluorescence assays and metabolomics. With the identity of bilirubin reductase in hand, we then surveyed thousands of gut metagenome samples from healthy adults, Inflammatory Bowel Disease (IBD) patients, and infants to assess the prevalence of bilirubin reductase in these populations. We found that bilirubin reductase was nearly universally present in healthy adults, while the prevalence of the gene was much lower in patients with IBD and in infants, especially during the first few months of life when infants are most susceptible to developing jaundice. Through the identification of bilirubin reductase, this study fills a long-standing knowledge gap in the heme degradation pathway and provides a new building block to understand disorders of serum bilirubin homeostasis like jaundice.

## Results

### Identification of an old yellow enzyme as a putative bilirubin reductase

Our goal was to pinpoint the gene encoding bilirubin reductase through experimental screening and comparative genomics. The bilirubin reductase enzyme is likely an oxidoreductase that acts on carbon-carbon double bonds (EC:1.3.-,-)(**Figure 1A, B**) and should be present in the genomes of species that can reduce bilirubin and absent in those that cannot. By examining closely related species with different abilities to reduce bilirubin, we aimed to identify a specific gene responsible for this activity.

**Figure 1:**
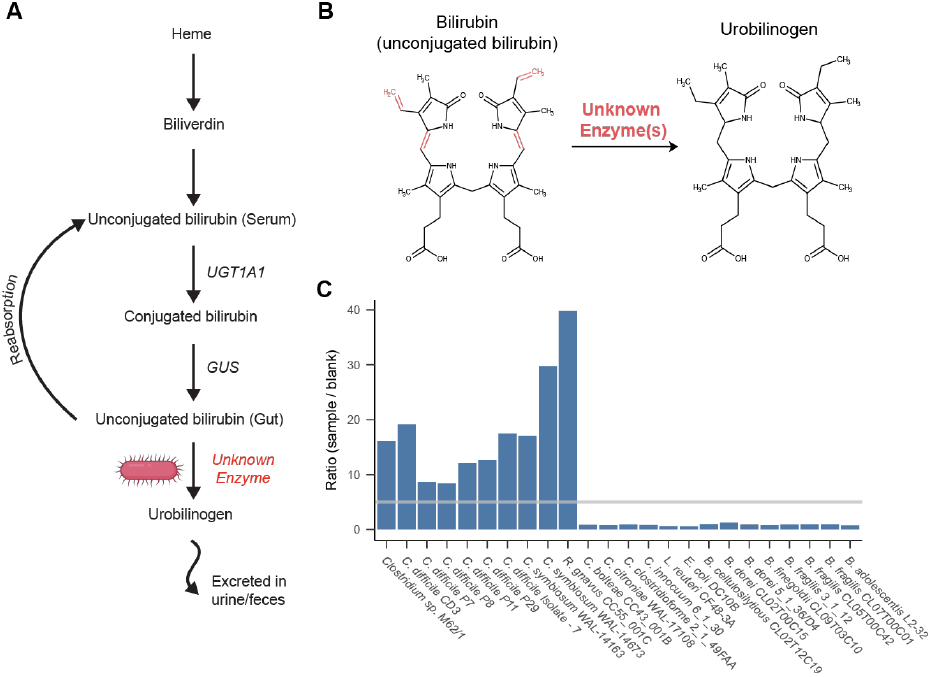
Identification of bilirubin reducing bacterial strains. A) Schematic representation of the heme degradation pathway. B) Diagram of the structures of bilirubin and urobilinogen. The bonds reduced during bilirubin reduction are shown in red. C) Results of fluorescence assay screening of bacterial strains. Bars show the ratio of the samples fluorescence to the corresponding experiments media blank. The gray line marks a ratio of 5, above which the sample was considered to be positive for bilirubin reduction.

To identify gut microbial species capable of bilirubin reduction, we employed a fluorescence assay on extracts from bacteria grown in media with bilirubin. The premise of the assay is that the unstable product of bilirubin reduction, urobilinogen, can be readily oxidized to the more stable urobilin, and measured via fluorescence (excitation 500 nm, emission 513 nm), while bilirubin itself is not fluorescent. The addition of iodine during the sample preparation for the fluorescence assay oxidizes all possible products of bilirubin reduction, including urobilinogen and its downstream product stercobilinogen, to urobilin and stercobilin. We considered a sample positive for bilirubin reduction if the magnitude of its fluorescence was greater than five times higher than the media blank. As a positive control, we demonstrated that the known bilirubin reducer, *C. difficile* CD3, returned a positive result in the fluorescence assay (**Figure 1C**)^25^. To confirm urobilin production by *C. difficile* CD3, we performed liquid chromatography with tandem mass spectrometry (LC-MS/MS) with standards for urobilin, the intermediate metabolite mesobilirubin, and stercobilin. Consistent with the fluorescence assay results, *C. difficile* media extracts contained ~8 times more urobilin than the media blank (Urobilin m/z 591.4→ m/z 343.2) while neither mesobilirubin nor stercobilin were detected (**Figure S1 A, B**). Thus, this fluorescence assay can be used as an effective screen for bacterial bilirubin reduction.

To search for additional bilirubin reducing species, we grew various bacteria from the major phyla of the human gut microbiome, prioritizing species previously reported to reduce bilirubin and their taxonomic relatives, in media supplemented with bilirubin and assayed for urobilin with fluorescence. Using a combination of fluorescence and LC-MS/MS we identified three species that were previously not known to be capable of bilirubin reduction: *Clostridium symbiosum* (strains WAL-14163 and WAL-14673), *Clostridium sp*. M62/1, and *Ruminococcus gnavus* CC55_001C (**Figure 1C**). Additionally, 14 gut species did not return a positive fluorescence result (**Figure 1C**). Of the tested species, all bilirubin reducers were found to be in the class Clostridia of the phylum Firmicutes. However, close relatives of these reducers, such as *Clostridium citroniae* WAL-17108, *Clostridium clostridioforme* 2_1_49FAA, and *Clostridium bolteae* CC43_001B, were not found to be reducers.

We leveraged the variability in bilirubin reduction between closely related strains to identify candidate bilirubin reductase genes through a comparative genomics analysis. Our analysis focused on the genomes of five confirmed bilirubin reducing strains and five closely related strains that did not show any bilirubin reduction activity in our fluorescence assay. Among the ten genomes analyzed, a total of 6,256 orthogroups were identified, of which 389 were predicted to be putative oxidoreductases. Only two of these orthogroups fit the same pattern of presence and absence in the strains as the bilirubin reduction phenotype. One of them was predicted to be a 4-Hydroxy-3-methylbut-2-enyl diphosphate reductase (EC: 1.17.1.2) which is an enzyme that acts on CH or CH2 groups and is involved in isoprenoid biosynthesis ^26^. Considering that the reaction mechanism of this enzyme and the metabolic pathway it is involved in showed little similarity to the expected mechanism of bilirubin reduction we looked further into the other candidate orthogroup. This other orthogroup was an unannotated enzyme and was found to be homologous to 2,4-Dienoyl CoA reductase (EC:1.3.1.34), an oxidoreductase that reduces carbon carbon double bonds, similar to the expected bilirubin reduction reaction. This enzyme was further analyzed as a putative bilirubin reductase.

We then analyzed the operons that contained the putative bilirubin reductases. Three versions of a putative bilirubin reductase operon were identified consisting of different combinations of three genes hereafter referred to as *bilQ, bilR*, and *bilS* (**Figure 2**). *bilR* is the putative bilirubin reductase, *bilS* is a flavodoxin-like protein, and *bilQ* is a MarR family transcriptional regulator. *C. difficile* CD3 and both strains of *C. symbiosum* contained versions of the operon that included all three genes. *Clostridium sp*. M62/1 had only *bilR* and *bilS*and *R. gnavus* CC55_001C had just *bilR*. The *R. gnavus* CC55_001C *bilR* differs in that it encodes two extra C-terminal domains in comparison to the other four *bilR* genes (**Figure S2**). The N-terminal domain of the *R. gnavus* CC55_001C BilR protein is predicted to be a TIM barrel fold, and it is homologous to the full protein sequences of other *bilR* genes. The two extra domains of the *R. gnavus* CC55_001C

**Figure 2:**
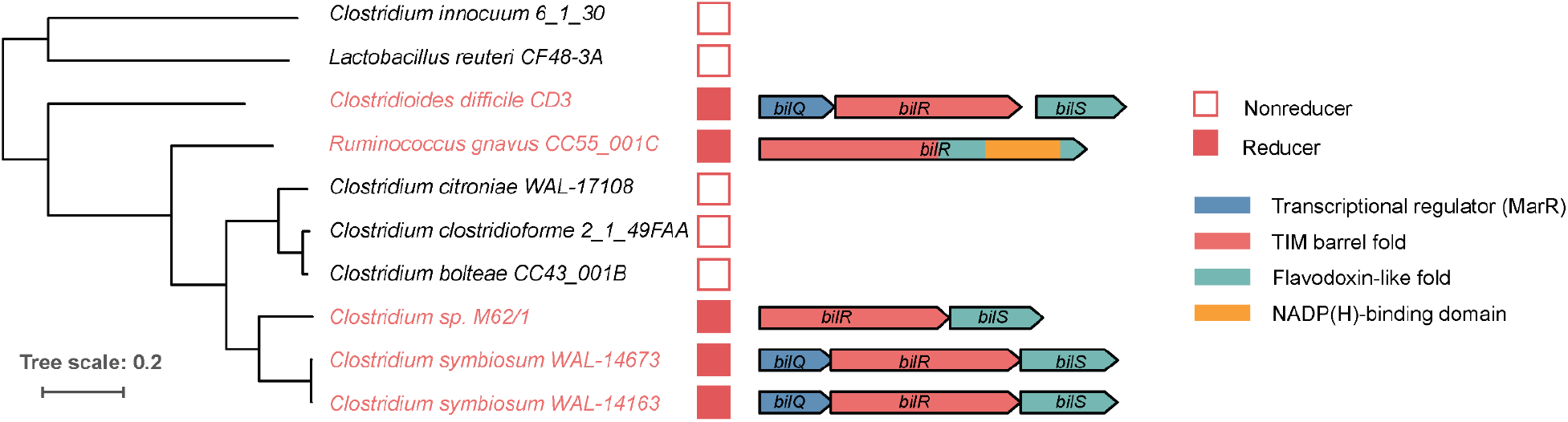
Putative bilirubin reductase operons. The phylogenetic tree shows the relationship between five bilirubin reducers and five non-reducers. Genes are represented as arrows. Genes are colored to show predicted domains.

BilR are a flavodoxin-like domain and an NADP(H)-binding domain. There was no sequence homology detected between the flavodoxin-like domain encoded by the *R. gnavus* CC55_001C *bilR* and the flavodoxin-like protein encoded by *bilS*, although they are predicted to have the same fold and are likely involved in electron transfer. The BilR of *R. gnavus* CC55_001C was predicted to belong to the Old Yellow Enzyme family (COG1902) and exhibits clear homology to *E. coli* 2,4-Dienoyl CoA reductase (PDB:1PS9)(protein identity=29.82%, RMSD of predicted BilR and 2,4-Dienoyl CoA reductase =2.79), a relatively promiscuous enzyme that catalyzes the reduction of both 2-trans, e-cis- and 2-trans, 4-trans-dienoyl-CoA thioesters ^27^. This homology supports the identification of BilR as a putative bilirubin reductase, as bilirubin reductase is hypothesized to be an oxidoreductase that can act on multiple CH-CH group donors of bilirubin.

### BilR confers bilirubin reductase activity

Once BilR was identified as a candidate bilirubin reductase, we sought to experimentally validate that it is sufficient to confer bilirubin reductase activity through ectopic expression in *E. coli*. The *bilRS* genes from *C. symbiosum* and *C. difficile* (short *bilRS)*, as well as *bilR* from *R. gnavus* (long *bilR)*, were separately cloned into a vector (pCW-lic) under the control of the inducible promoter *tac* and then transformed into *E. coli* 10-beta (**Figure 3 A, B**). Then, we induced the expression of *bilR or bilRS* with isopropyl β-D-1-thiogalactopyranoside (IPTG) and incubated the transformed bacteria with bilirubin. After 24 hours, we assayed for the reduction of bilirubin to urobilinogen via both a fluorescence assay and LC MS/MS. Both the long BilR and short BilRS reduced bilirubin to urobilinogen (fluorescence assay: *C. difficile* P=9.99e-06, *C. symbiosum* P=2.29e-05, *R. gnavus* P=1.01e-4, two sample t-test on log2 transformed data; metabolomics: *C. difficile* P=7.59e-05, *C. symbiosum* P=2.78e-04, two sample t-test on log2 transformed data (urobilin m/z 591.4→ m/z 343.2)) (**Figure 3 C, D, E, F**). *E. coli* with the empty vector backbone pCW-lic, which was used as a negative control, did not reduce bilirubin

**Figure 3:**
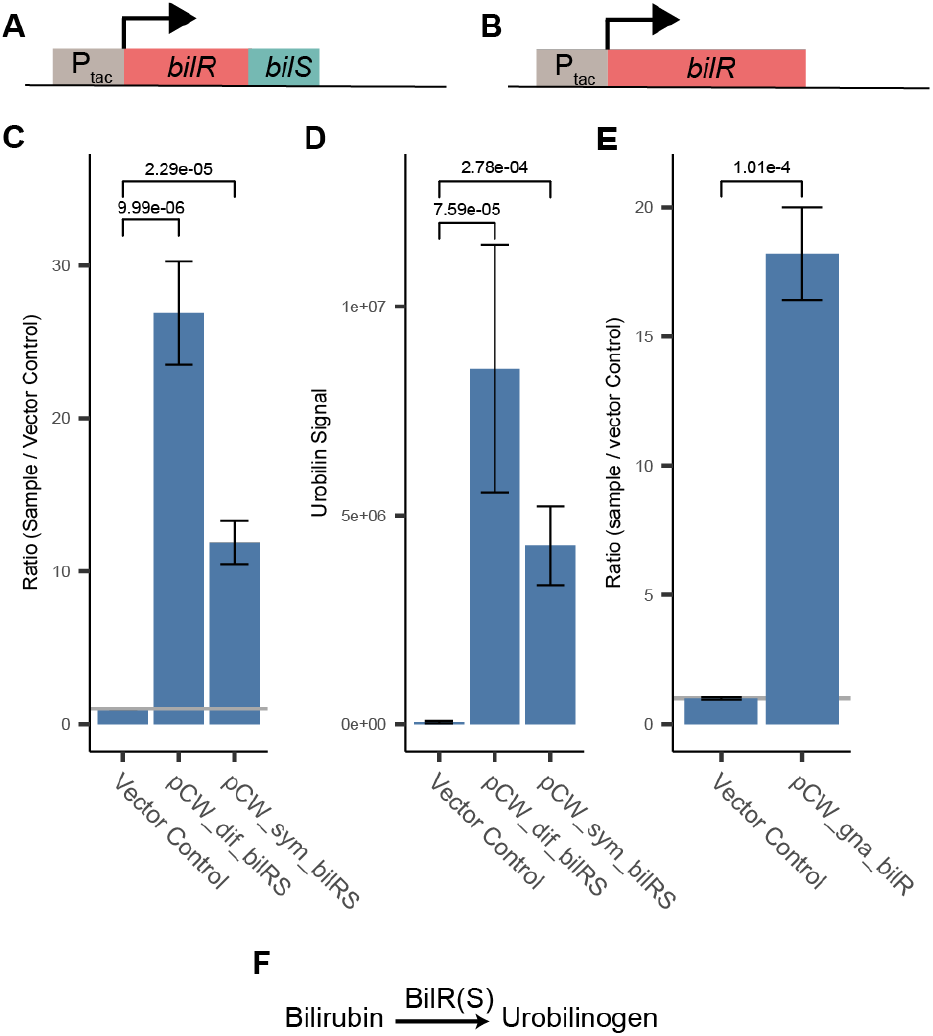
Confirmation of *bilR(S)* bilirubin reductase activity. A) Schematic of *C. symbiosum* and *C. difficile bilR(S)* construct. B) Schematic of *R. gnavus bilR* construct. C) Fluorescence assay comparing bilirubin reduction activities of *E. coli* 10-beta transformed with empty vector, *C. symbiosum* construct, and *C. difficile* construct (two-sample t-test). D) Metabolomics confirmation of urobilin production (two-sample t-test). E) Fluorescence confirmation of bilirubin reduction activity of *E. coli* 10-beta transformed with the *R. gnavus bilR* (two-sample t-test). F) Representation of the reaction catalyzed by the bilirubin reductase enzyme. Bar heights in plots indicate the mean values of biological quadruplicates (C and D) or triplicates (E). Error bars indicate one standard error above and below the mean values. P-values of a two sample t-test testing if the mean of the samples is greater than the mean of the vector controls are provided above the brackets.

(P=0.167, one-sided t-test on log2 transformed data). Neither mesobilirubin (m/z 589.3 →m/z 301.2) nor stercobilin (m/z 595.4→m/z 345.2) were consistently detected (**Figure S3**). To validate that bilirubin reductase is capable of all four carbon-carbon double bond reductions, we also incubated *E. coli* expressing *bilRS* with mesobilirubin, an intermediate metabolite of the proposed reaction, and showed that BilRS reduced mesobilirubin to urobilinogen (**Figure S1 C).** These results support that BilR is sufficient to reduce bilirubin to urobilinogen.

### Delineation of the BilR clade

To delineate the BilR sequences from other members of the Old Yellow Enzyme family, a combination of ancestral sequence reconstruction and structural analysis was used to identify key residues that were conserved within putative BilR sequences and that differentiated BilR from the other sequences (**Figure 4A**). Our analysis has revealed that one particular clade, referred to as Clade 1, is likely to be the bilirubin reductase clade (**Figure 4A**). This inference is supported by the fact that the predicted substrate binding pocket of sequences within Clade 1 is highly conserved, indicating that the specificity for bilirubin binding has been maintained (**Figure 4B**). Additionally, the branch connecting the parent node (Node 1) of Clade 1 and its direct ancestor (Node 2) is the most recent branch where changes in the residues of the substrate binding pocket of BilR occur (**Figure 4A, 4B, Figure S4 and S5**). These changes include mutations in a region that corresponds to a catalytic residue (Tyr-166) of the *E. coli* 2,4-dienoyl-CoA reductase. Two residues in the region, D166 and R167, were nearly universally conserved within Clade 1, and were predicted to be mutated from “XHGY” based on ancestral sequence reconstruction (**Figure 4C**). Changes in the catalytic machinery of enzymes frequently result in functional shifts, and this likely marks the point of divergence of bilirubin reductase from other members of the Old Yellow Enzyme family.

**Figure 4:**
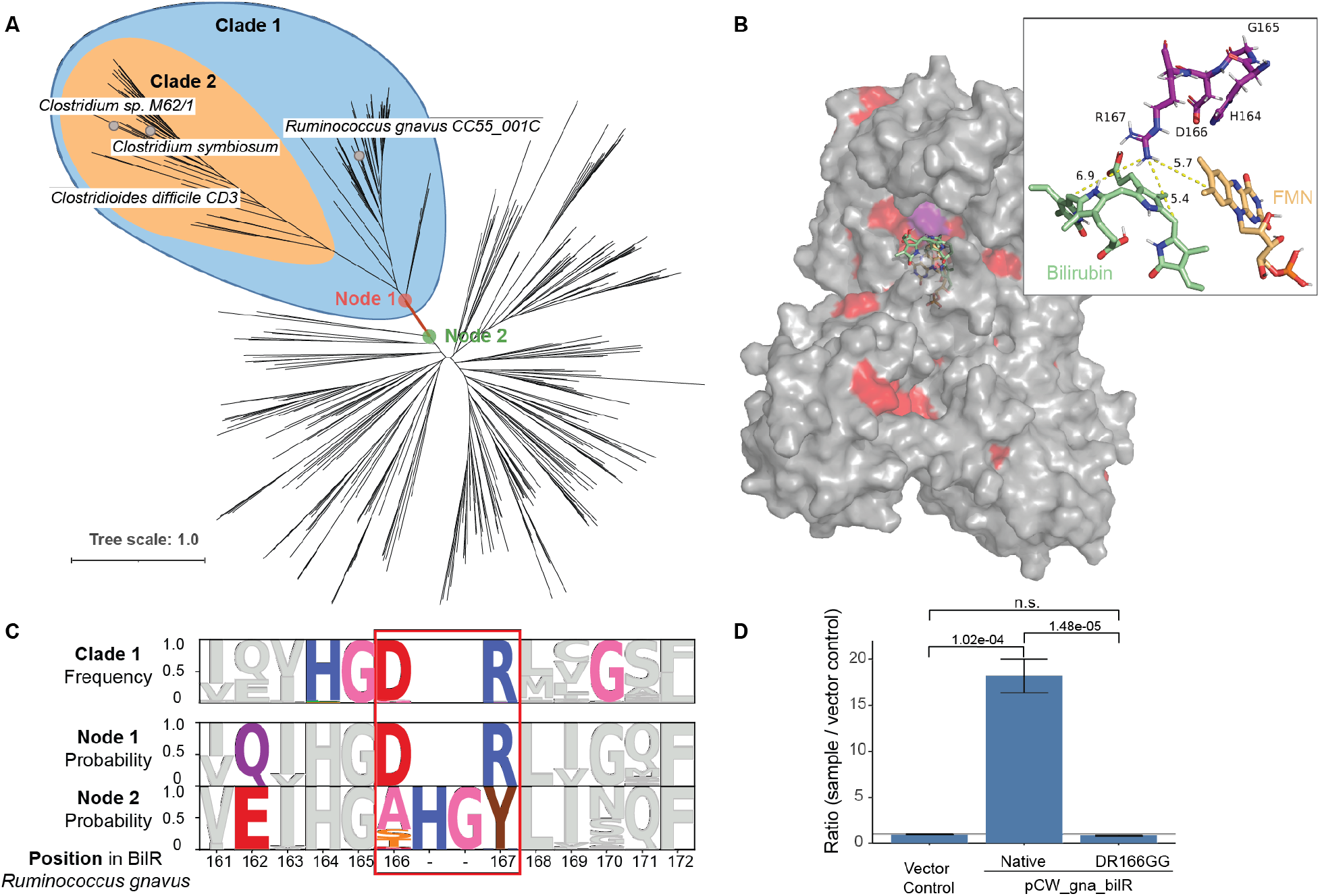
Identification of a *bilR* clade. A) Gene tree constructed from putative *bilR* sequences and related reductases from the Old Yellow Enzyme family. Experimentally confirmed bilirubin reducers are labeled in the tree. Clade 1 indicates the overall *bilR* clade, while Clade 2 indicates the short *bilR* clade. B) Predicted structure of the *R. gnavus* BilR with sequences conserved in greater than 90% of the BilR sequences colored as red. The HGDR residues are colored in purple. A docked bilirubin (green) and FMN molecule (orange) are shown on the structure and the inset shows the potential site of interaction between the R167 residue and the putative enzyme substrates. C) Diagram showing the conservation of each position within the BilR sequences (top), and predicted likelihood of each amino acid based on the ancestral sequence reconstruction for the BilR common ancestor (middle) and ancestor of other sequences in the tree (bottom). Positions are colored to highlight positions that differed between Node 1 and Node 2, positions that changed and were conserved within the BilR clade, and the HGDR motif identified in BilR. D) Fluorescence assay results comparing bilirubin reductase active in *E. coli* 10-beta transformed with a vector control, *R. gnavus bilR*, and *R. gnavus bilR* with the DR residues at position 166-167 mutated to GG. Bar heights in plots indicate the mean values of biological triplicates. Error bars indicate one standard error above and below the mean values. P-values of a two sample t-test testing if the mean of the native *R. gnavus bilR* was higher than the mean of the vector control or mutant.

To confirm that Clade 1 is indeed composed of bilirubin reductase enzymes, we performed experiments to investigate the functional impact of the D166R167 residues on the ability of the enzyme to reduce bilirubin. This included mutating the residues in question to G166G167 and ectopically expressing the mutated protein to measure the impact on the enzyme’s activity. The mutated protein demonstrated a significantly lower bilirubin reductase activity (P=1.48e-05, two sample t-test), showing a complete loss of functionality (**Figure 4D**). This experimental approach provides direct evidence that the “DR” residues in the active site are critical for bilirubin reduction, a characteristic that is specific to bilirubin reductase enzymes and distinct from other members of the Old Yellow Enzyme family. Combined with structural data, this supports the conclusion that Clade 1 is indeed the bilirubin reductase clade.

### The BilR bilirubin reductase is predominantly encoded by gut Firmicutes

With the ability to distinguish BilR from other reductases, we studied the taxonomic distribution and diversity of bilirubin reducers. We searched for *bilR* gene in the representative genomes of the Genome Taxonomy Database (GTDB) and identified putative bilirubin reductases in 658 species (**Figure 5**). The distribution of *bilR* genes is largely in agreement with previously reported bilirubin-reducing species. The representative genomes of *C. ramosum* and *C. difficile* were predicted to contain *bilR* genes. While the representative genome of *C. perfringens* does not have bilirubin reductase, 137 out of 315 genomes assigned to *C. perfringens* have it, supporting that bilirubin reduction is a strain-specific feature. The only inconsistency is that no species within the *Bacteroidaceae* family were predicted to be bilirubin reducers, but a study conducted in 1972 showed that *Bacteriodes fragilis* can reduce bilirubin ^8^. This study did not provide strain information and the genome is unavailable, making it possible that the observed bilirubin reduction is a strain-specific feature or due to an incorrect assignment of taxonomy.

**Figure 5:**
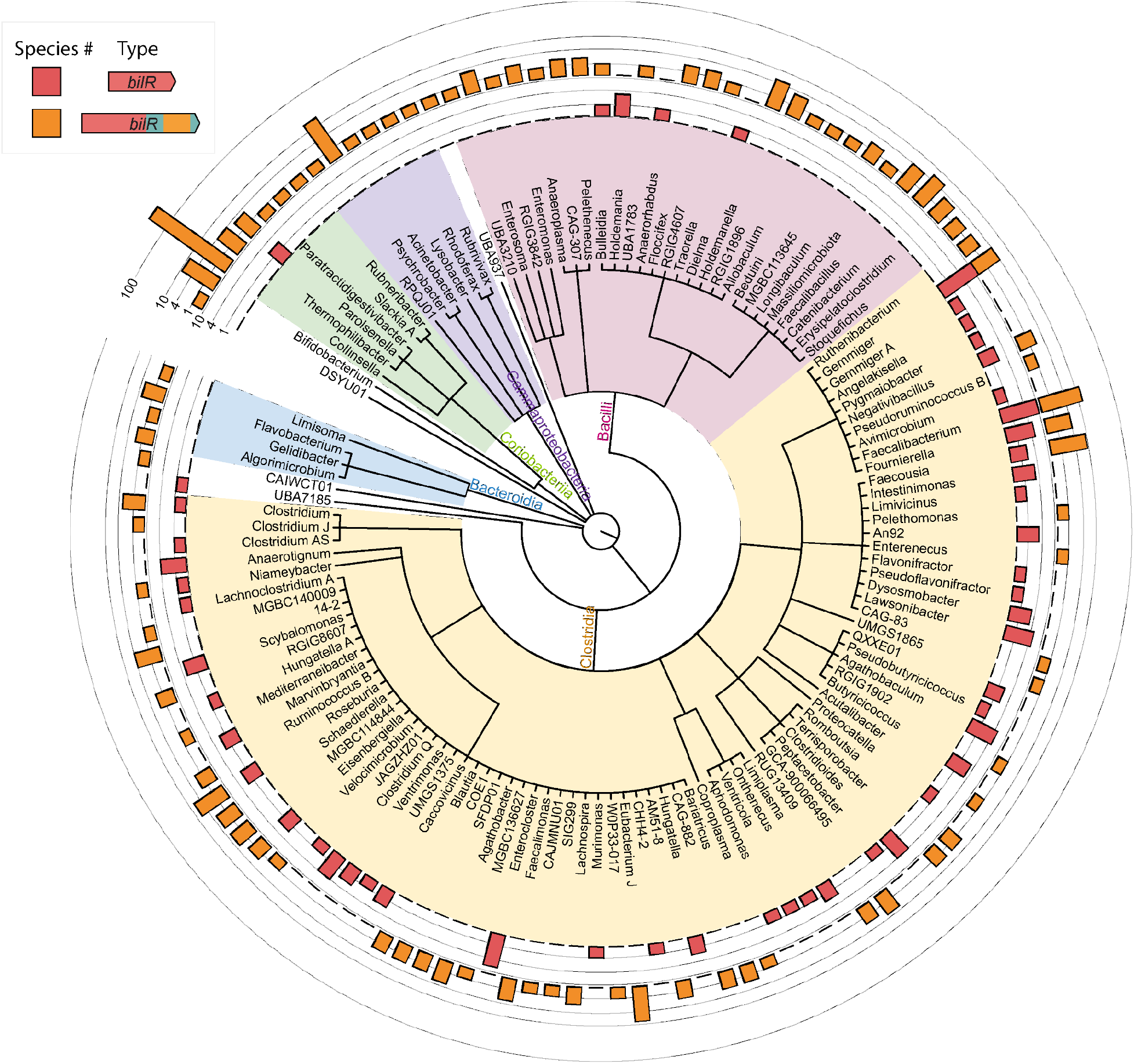
Taxonomic distribution of bilirubin reductase. The cladogram shows the relationships between different taxa with detected bilirubin reductase genes. The outer rings show the presence of the short *bilR* gene (red) and long *bilR* gene (orange).

The long variant of the *bilR* gene is widely distributed across multiple phyla, while the short variant seems to be more exclusive to Firmicutes, only being seen in the *Parolsenella* genus outside of Firmicutes. The majority of *bilR* genes were identified in Firmicutes species, with a mixture of both versions being seen. The short variant of the *bilR* gene is monophyletic (Clade 2 in **Figure 4A**), suggesting that there has been extensive horizontal gene transfer during the evolution of *bilR*, particularly in Firmicutes. A significant proportion of Firmicutes bilirubin reducers have been isolated from the human gut. This includes common human gut bacteria species such as *Roseburia intestinalis, Roseburia inulinivorans*, and *Faecalibacterium prausnitzii*. In contrast, bilirubin reducers identified in the Bacteroidota phylum were mostly limited to the order *Flavobacteriales*, which are typically found in aquatic and soil environments. Nine species in the genus *Bifidobacterium* were predicted to be bilirubin reducers and were isolated from the feces of non-human mammals such as chickens, lemurs, and monkeys. This implies that the bilirubin reducing capacity of the human gut microbiota is largely influenced by the abundance of Firmicutes-associated bilirubin reducers.

### The BilR bilirubin reductase is frequently absent in the neonate gut microbiome

To examine the relationship of bilirubin reductase to age and health we performed a large scale analysis of human gut metagenomes. Infants are known to be susceptible to hyperbilirubinemia and jaundice, especially within the first few days-to-weeks of life^28^. We examined the presence and absence of bilirubin reductase across 4296 infant gut metagenomes from the first year of life. In the samples analyzed we saw a stark trend where 78% of samples taken from within the first month did not have bilirubin reductase present but by the end of the first year of life only 5% of samples were missing bilirubin reductase (**Figure 6A**). There is a high level of bilirubin reductase absence over the first few months of life, corresponding to the general period of time where neonatal jaundice risk is highest (**Figure S6**). The bilirubin reductase presence also appears to stabilize relatively quickly with the presence becoming relatively stable after around 6 months, with the largest changes in bilirubin reductase presence being seen between within the first 180 days of life. The correspondence between the period of life where we observe the most samples missing bilirubin reductase and the risk of neonatal jaundice suggests a strong connection between the microbiome composition and the development of jaundice in infants.

**Figure 6:**
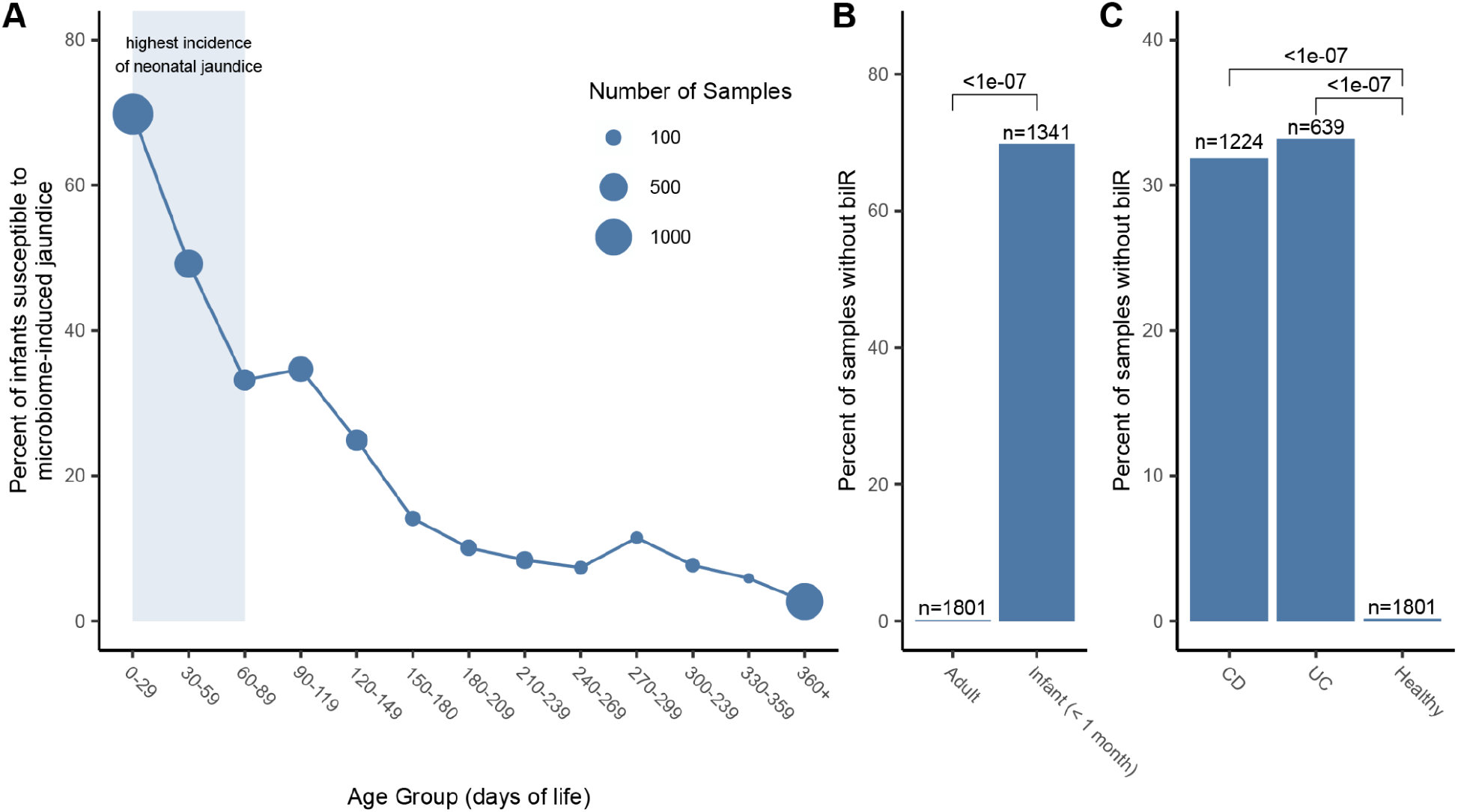
Presence of *bilR* in the human gut during development and disease. A) Percentage of infant gut metagenomes missing *bilR* during their first year of life. The period of highest jaundice susceptibility is indicated by a shaded blue area on the plot. B) Comparison of *bilR* absence in samples from healthy adults and infants in their first month of life. C) Comparison of the percentage of samples with no *bilR* detected from healthy adults and adults with IBD. The number of metagenomic samples included in each dataset is indicated above each bar. The P-values for each comparison show the results of a test of equal proportions testing if the fraction of samples with no *bilR* detected was different between groups.

### Bilirubin reduction by BilR is a core function of the healthy human gut microbiome that is frequently absent in IBD patients

When examining metagenomes from healthy adults we found that bilirubin reductase was absent in only 0.1% of the 1801 samples analyzed, a significantly lower fraction of samples (P<2.2e-16, test of equal proportions) than was seen in the infants during the period of highest jaundice susceptibility (**Figure 6B**). While bilirubin appeared to be nearly ubiquitous in healthy adults, IBD patients have been observed to have significantly lower levels of urobilin^29^, the product of bilirubin reduction, and have been observed to form pigmented gallstones which are gallstones containing high levels of bilirubin^30–32^. We compared the bilirubin presence in bilirubin reductase in 1863 gut metagenomes from IBD patients to the healthy samples, finding that bilirubin reductase was absent in a significantly higher fraction of samples from patients with Crohn’s disease (P<2.2e-16, test of equal proportions) or ulcerative colitis (P<2.2e-16, test of equal proportions) (**Figure 6C**). This provides evidence that the microbial dysbiosis that is often associated with IBD and other inflammatory diseases may be having a direct impact on the bilirubin reduction activity of the gut microbiome.

## Discussion

Despite the identification of urobilin as the yellow pigment in urine more than 125 years ago, the enzymes responsible for its production have remained a mystery^33^. This knowledge gap has hindered our understanding of how bilirubin reduction in the gut affects serum bilirubin levels and the role it plays more generally in human health and disease. Here, through a combination of biochemical analyses and comparative genomics, we have identified and characterized bilirubin reductase, an enzyme responsible for the reduction of bilirubin to urobilinogen. Though it was previously thought that multiple enzymes were involved in the reduction of bilirubin to urobilinogen, our results support that a single enzyme performs the reduction of bilirubin to urobilinogen^25^. The identification of bilirubin reductase allowed us to profile its abundance in 7,960 metagenomes showing that bilirubin reduction is a core feature of a healthy adult human microbiome^34,35^. In addition, we found decreased prevalence of bilirubin reductase in neonates during the period of the highest incidence of neonatal jaundice. Furthermore, we found that the prevalence of bilirubin reductase is decreased in IBD patients.

Bilirubin reduction is the key step that determines the degree to which the heme degradation byproduct bilirubin is reabsorbed or excreted^13^. Bilirubin is insoluble and can easily be reabsorbed into circulation, but urobilinogen and stercobilinogen are water-soluble and readily excreted in urine and feces preventing reabsorption and promoting excretion of bilirubin. The importance of this process is further highlighted by the limited capacity of the UGT1A1 enzyme in the liver to conjugate serum bilirubin which has been shown to be a rate limiting step in heme degradation^36–39^.

While the discovery of bilirubin reductase illuminates a key gap in our understanding of the health relevant steps of heme excretion, there is an additional downstream step performed by gut microbes with an unexplained mechanism: the reduction of urobilinogen into stercobilinogen. Similar to bilirubin, urobilinogen may serve as a terminal electron acceptor for microbes encoding these reductases, providing them a competitive advantage in the anaerobic environment of the gut. However, once bilirubin is reduced to urobilinogen it is already destined for excretion in urine or feces and further reduction to stercobilinogen does not change its fate^40,41^.

With the knowledge of the species, genes, and enzymes involved in bilirubin reduction, future research can now focus on to what extent gut microbial bilirubin metabolism affects serum bilirubin homeostasis and the role of bilirubin reduction in health and disease. It has been hypothesized that neonatal jaundice may be exacerbated by the absence or low abundance of bilirubin reducing microbes in the gut^14^. Our results support this hypothesis and show that the period in which neonates experience the highest incidence of jaundice corresponds with the lowest prevalence of bilirubin reducing microbes. In order to further investigate this hypothesis cohort studies that simultaneously measure serum bilirubin, fecal urobilinoids, and the absolute abundance of bilirubin reducing bacteria in infants are needed. Additionally, the lowered prevalence of bilirubin reducing bacteria in IBD patients leads us to hypothesize that the combination of excess unconjugated primary bile acids and unconjugated bilirubin specifically in IBD patients could contribute to the increased incidence of calcium bilirubinate gallstones noted in these patients^30–32^. Our work fills in a long standing knowledge gap, providing foundational knowledge that can serve as the basis of the further investigation of the importance of bilirubin metabolism in human health.

## Materials and Methods

### Culture methods

#### Anaerobic bacteria

Bacterial strains were obtained from the NIH Biodefense and Emerging Infections Research Resources Repository (BEI). Strains were inoculated from a glycerol stock and grown under anaerobic conditions (90% N_2_, 5% CO_2_, 5% H_2_) at 37°C in an anaerobic chamber (Coy Laboratory Products). Strains were grown in 200 mL of liquid brain heart infusion broth (BHI, Research Products International (RPI), B11000) or yeast casitone fatty acid broth (YCFA) with 4.4 mg/100 mL sterile filtered bilirubin dissolved in dimethylsulfoxide (DMSO, Sigma-Aldrich). The bacteria were grown anaerobically for 24 hours before starting the chloroform extraction.

#### Transformed E. coli

Transformed *E. coli* strains were inoculated from an agar colony plate into 300 mL BHI media with 100 ug/mL carbenicillin (GoldBio, 00901C103) and 100 uM isopropyl β-d-1-thiogalactopyranoside (IPTG, GoldBio, 221105I2481) and shaken aerobically at 37°C for 16 hours. The bacteria were then pelleted by centrifugation at 3260 RCF for 6 minutes and moved to the anaerobic chamber. The pellet was resuspended in 200 mL of BHI with 4.4 mg/100 mL sterile filtered bilirubin dissolved in DMSO, 100 ug/mL carbenicillin, and 100 uM IPTG to induce expression of the cloned *bilRS*. The strains were incubated anaerobically for 24 hours before starting the chloroform extraction.

### Measuring bilirubin reduction

#### Fluorescence Assay

Bacterial cultures were removed from the anaerobic chamber and pelleted by centrifugation at 3260 RCF for six minutes. The supernatant was filtered through a non-sterile 0.22 um filter. Five mL of chloroform (Sigma-Aldrich) was added to the filtered supernatant and separated via organic extraction in a separatory funnel. The resultant chloroform-urobilinogen solution was air dried until the chloroform was completely evaporated, producing the extract used in the subsequent fluorescence assays and LC MS/MS.

Because the urobilinogen product of bilirubin reduction is unstable in oxygen, measuring urobilinogen itself was infeasible. Therefore, to measure bilirubin reduction, urobilinogen was oxidized to urobilin via the addition of iodine. The urobilin was quantified by adding zinc acetate solution to form a fluorescent zinc-urobilin complex.^42–44^ However, this assay also detects stercobilin, which fluoresces at the same wavelength as urobilin.^45,46^ To differentiate the two potential products, the fluorescence assay was verified by mass spectrometry.

The instability of urobilinogen requires this assay be performed immediately after the chloroform from the chloroform extraction had evaporated. The extracts were rehydrated using 600 μL of deionized water. A 400 μL aliquot of the rehydrated solution was transferred to a separate tube to continue the assay. 10 μL of 10% Povidone-iodine solution (CVS, A47194) was added to oxidize the urobilinogen to urobilin, followed by 10 μL of a 100 mM cysteine solution to reduce the rest of the iodine to prevent bilirubin oxidation after addition of Schlesinger’s reagent (545 mM zinc acetate in methanol). 400 μL of Schlesinger’s reagent was added to increase the fluorescence of urobilin. The resulting solution was portioned into 100 μL triplicates or quadruplets onto a 96 well acrylic plate. Each well’s fluorescence was measured by SpectraMax M5 plate reader using a 495/525 nm wavelength at medium gain. Samples that produced a fluorescence signal greater than five times the fluorescence of the media blank were identified as reducers. Samples below the five times threshold were considered to be non-reducers.

#### LC with tandem mass spectrometry (LC-MS/MS)

1 mL of the chloroform solution from the chloroform extraction was aliquot and allowed to dry completely. The samples were sent with overnight shipping in dry ice to the Duke metabolomics core. 60 uL of 10% Povidone-iodine solution was added to the dried samples. Samples were vortexed, then centrifuged at 15,000 rpm for 5 minutes at 4°C. The supernatant was transferred into the 1.7 mL Macherey-Nagal glass vial for the LC-MS/MS injection.

Samples were analyzed with the 6500+ QTRAP LC-MS/MS system (Sciex, Framingham, MA). The Sciex ExionLC UPLC system includes a degasser, an AD autosampler, an AD column oven, a controller, and an AD pump. The liquid chromatography separation was performed on an Agilent Eclipse Plus C18 RRHD column (2.1 x 50mm, 1.8 μm) with mobile phase A (0.2% formic acid in water) and mobile phase B (0.2% formic acid in acetonitrile). The flow rate was 0.35 mL/min. The linear gradient was as follows: 0-0.5min, 100% A; 4.0-5.5min, 0%A; 5.6-7.5min, 100%A. The autosampler was set at 10°C and the column was kept at 35°C. The injection volume was 5 μL. Mass spectra were acquired under positive electrospray ionization with the ion spray voltage of 5500 V. The source temperature was 450 °C. The curtain gas, ion source gas 1, and ion source gas 2 were 33, 55, and 60 psi, respectively. Multiple reaction monitoring (MRM) was used to detect Mesobilirubin (m/z 589.3 →m/z 301.2) Urobilin (m/z 591.4→ m/z 343.2) and Stercobilin (m/z 595.4→m/z 345.2). All data was analyzed in software Analyst 1.7.1.

### Construct development

#### pCW-sym-bilRS

To ectopically express the *bilRS* genes from *C. symbiosum* in *E. coli*, the gene was amplified and inserted into an expression vector backbone pCW-lic (Addgene, 26908). A polymerase chain reaction (PCR) amplification using OneTaq Master Mix (NEB, M0482) was performed on *C. symbiosum* genomic DNA with forward primer TAAGCACATATGCTGGGAAAGACGAAGGAGGCTC and reverse primer TAAGCAGGTACCGGCAGCGTCCCTGAGT to amplify *C. symbiosum bilRS*. The product was purified with a Monarch PCR and DNA cleanup kit (NEB, T1030) and restricted with enzymes Nde1 (NEB, R0111) and Kpn1-HF (NEB, R3142) using NEBcloner® protocols to prepare for ligation into the pCW-lic backbone. The pCW-lic vector backbone was restricted with the same enzymes. pCW-lic was dephosphorylated with antarctic phosphatase (NEB, M0289) to prevent self-ligation. The restricted insert and dephosphorylated backbone were ligated with T4 DNA ligase (NEB, M0202) to form the construct.

#### pCW-dif-bilRS

The same protocol was used as for pCW-sym-bilRS, with two exceptions. PCR amplification was performed on *C. difficile* genomic DNA with forward primer TAAGCAGGATCCCAGCTGTGGAGGAAGAATAGGATG and reverse primer TGCTTAAAGCTTCTACTACACAATACTAGCTTTAATCATCATA to amplify *C. difficile bilRS*. The product was restricted with enzymes BamH1-HF (NEB, R3136) and HindIII-HF (NEB, R3104) before ligating.

#### pCW-gna-bilR

The same protocol was used as for pCW-sym-bilRS, with two exceptions. PCR amplification was performed on *R. gnavus* genomic DNA with forward primer TAAGCAGGATCCGCAGAAAGAAAATGTTAAGGAGAGGCTG and reverse primer TAAGCAGGTACCGAAGGATGTTTCATCCACCTGTACG to amplify *R. gnavus bilR*. The product was restricted with enzymes BamH1-HF and Kpn1-HF before ligating.

#### pCW-gna-bilR(DR166GG)

A PCR amplification using Phusion® High Fidelity DNA Polymerase (NEB, M0530S) was performed on pCW-gna-bilRS with forward primer GAAGTACACGGAGGGGGGCTGATCGGATCATTCTGTTCG and reverse primer GAATGATCCGATCAGCCCCCCTCCGTGTACTTCTATAATATC to amplify the pCW-gna-bilRS plasmid and introduce a mutation of the 166 aspartic acid and 167 arginine to glycines. The product was restricted with Dpn1 (NEB, R0176S) to remove the template DNA and purified with a Monarch PCR and DNA cleanup kit before transforming.

#### Transformation

The constructs were individually transformed into NEB 10β competent *E. coli* cells using the manufacturer’s protocol (NEB, C3019). The cells were plated on Luria-Bertani (LB) plates with 100 ug/mL carbenicillin to select for successfully transformed colonies. The transformed colonies were verified through Sanger sequencing by Azenta.

### Bioinformatic Analysis

#### Identification of candidate bilR genes

Five genomes from the experimentally confirmed bilirubin reducers and five genomes from closely related confirmed non-reducers were downloaded from NCBI. The assembled genomes for *C. bolteae* CC43_001B and *Clostridium innocuum* 6_1_30 were not available so the raw sequencing data was downloaded from their corresponding bioprojects. Fastq files for the two species were download from SRA using SRA-Tools (version 2.11.0)(https://github.com/ncbi/sra-tools), trim-galore (version 0.6.7) (https://github.com/ncbi/sra-tools) was used to trim and quality filter the fastq files, and the genomes were assembled using spades (version 3.15.5) ^49^. The genomes were annotated with Prokka (version 1.14.6) ^50^. Orthologer (version 2.7.1) was used to group the predicted protein sequences from each genome into orthogroups using default settings ^51^. ECPred (version 1.1) was then used to assign the putative E.C. numbers to each protein sequence using default setting ^52^. The orthogroups were then subset to only keep groups that contained more than 2 proteins assigned to the oxidoreductase E.C. number (EC: 1.-,-,-) leaving a total of 389 orthogroups. The taxonomic distribution of the proteins within these oxidoreductase orthogroups was profiled to identify orthogroups which were present in the bilirubin reducing bacteria and absent in the nonreducing species. Only two orthogroups fit the taxonomic distribution. Of them, one was assigned to EC:1.17.7.4 (4-hydroxy-3-methylbut-2-enyl diphosphate reductase) and unlikely to be bilirubin reductase; the other one was assigned to EC:1.-,-,- and was further investigated as a putative bilirubin reductase.

#### Structural prediction and molecular docking

The structures for the putative BilR protein from *R. gnavus* CC55_001C and the BilR and BilS proteins from *C. symbiosum* WAL-14163 were predicted using AlphaFold (version 2.2.0) ^53^. Putative substrate binding pockets were predicted using Fpocket (version 4.0.2) with default settings ^54^. The predicted pockets were visualized using PyMOL (version 2.5.0) and were compared to the substrate binding region of the E. coli 2,4-Dienoyl CoA reductase (PDB:1PS9) to identify three putative substrate binding regions on the BilR structure (http://www.pymol.org/). Structures for bilirubin (PubCham CID 5280352) and FMN (PubChem CID 643976) were docked on the R. gnavus BilR structure using AutoDock Vina (version 1.2.0) ^55,56^. The docking simulations were performed within 20 by 20 by 20 angstrom cubes centered on the centerpoints of the three chosen Fpocket substrate binding pocket predictions with the exhaustiveness set to 32. The docking results were visualized using PyMOL and the results using within figures were chosen based on what is known about what bonds are reduced on bilirubin and based on comparisons to the known binding conformations of the E. coli 1PS9 protein. Protein structural alignments were performed between the *Ruminococcus gnavus* CC55_001C predicted structure and each of the *Clostridium symbiosum* WAL-14163 BilR, *C. symbiosum* WAL-14163 BilS, and *Escherichia coli* 1PS9 structures using TM-Align ^57^.

#### Ancestral sequence reconstruction

All protein sequences from the GTDB representative genomes (release 207) assigned to COG1902 (Old Yellow Enzyme family) by eggNOGmapper (version 2.1.6) were clustered at 70% identity using CD-HIT (version 4.6.8) ^58–60^. A search was performed using DIAMOND (version 2.0.9) with the five putative BilR sequences as queries and the results were filtered to only keep hits that had less than 1e-48 e-value and greater than 30% identity to at least one of the query sequences ^61^. The sequence for the *R. gnavus* CC55_001C BilR was added to this set of proteins and was aligned using Clustal Omega (version 1.2.4) ^62^. The alignment was trimmed to remove positions that have more than 97% of gaps. A phylogenetic reconstruction was performed based on the trimmed alignment using IQ-TREE 2 (version 2.1.2) ^63^. An ancestral sequence reconstruction was then performed using GRASP (version 30-July-2022) with default settings for both the joint and marginal reconstructions ^64^.

#### Search for BilR in GTDB database

The alignment of the amino acid sequences from species in Clade 1 (as shown in **Figure 4A**) and the positions in the alignment corresponding to the first 373 residues of the *R. gnavus* CC55_001C BilR protein were extracted to generate an HMM profile using the hmmbuild tool. This profile represents the TIM barrel domain of the BilR proteins. The HMM profile was then used to perform a search within genomes of the GTDB ^60^ to profile the taxonomic distribution of BilR. The search was performed using hmmsearch and only hits below an e-value of 1e-100 were considered. A protein domain prediction was then performed on the putative BilR hits using InterProScan (version 5.57-90.0) ^65^. The putative BilR sequences were filtered requiring that they be at least 50% of the length of the BilR sequences from *C. symbiosum* WAL-14163, they have the highly conserved HGDR motif present in their sequence, and that they only have the expected domains present (PF00724 for the shorter bilR or PF07992 and PF00724 for longer BilR version). The presence or absence of BilR was then summarized across the different bacterial taxa and the results were visualized using iTOL ^66^.

#### Profiling of bilR presence in the human gut

A collection of metagenome datasets passing basic quality control was curated to provide a cross section of gut metagenomes from infants in the first year of life (n=4296), IBD patients (n=1863), and healthy adults (n=1801). A *bilR* reference gene dataset was generated by searching the Unified Human Gastrointestinal Genome collection using the previously generated HMM profile of the BilR TIM barrel domain. The resulting hits were filtered based on a 1e-100 e-value threshold and were subjected to the same quality control used in the search against the GTDB database, resulting in a total of 11158 *bilR* reference sequences. The metagenomes were all processed using a standard workflow that consisted of the following steps 1) The reads were downloaded from SRA, 2) the adaptors were trimmed using Trim Galore with default settings, 3) potential human contaminant reads were identified by mapping the reads to a human genome reference (assembly T2T-CHM13v2.0) and were removed using Samtools (version 1.16.1) ^67^, 4) samples with less than one million reads were not kept, 5) reads were aligned the *bilR* reference gene set using Bowtie2 ^68^. The number of reads mapped to the *bilR* reference database was then summarized for each sample by normalizing by the total number of reads in the sample and multiplying by one million to give a *bilR* counts per million value (CPM).

Infant related metagenomes were binned into age groups of 30 days from 0 days to 1 year of life and samples without age metadata were not considered in this analysis. Samples from IBD patients were categorized based on if the patient had Ulcerative Colitis or Crohn’s Disease. For healthy gut metagenomes samples were excluded if they were from patients under three years of age. *bilR* was considered to be present in a sample if the *bilR* CPM value was greater than five CPM. The prevalence of *bilR* between groups was compared using a test of equal proportions, using the *prop.test* function from the R *stats* library, to test if the given proportion of samples with *bilR* present was different between two groups.

## Acknowledgements

This work utilized the computational resources of the NIH HPC Biowulf cluster. (http://hpc.nih.gov). pCW-LIC was a gift from Cheryl Arrowsmith (Addgene plasmid # 26098; http://n2t.net/addgene:26098; RRID:Addgene_26098). UPLC analyses of bilins were performed at the Iron and Heme Core facility at the University of Utah, supported in part by a grant from the NIH National Institute of Diabetes and Digestive and Kidney Diseases, Grant number U54DK110858. The following reagents were obtained through BEI Resources, NIAID, NIH: Clostridium symbiosum, Strain WAL-14163, HM-309; Ruminococcus gnavus, Strain CC55_001C, HM-1056; Clostridium clostridioforme, Strain 2_1_49FAA, HM-306; Clostridium bolteae, Strain CC43_001B, HM-1038; Escherichia coli, Strain DC10B, HM-49804; Lactobacillus reuteri, Strain CF48-3A, HM-102; Bacteroides finegoldii, Strain CL09T03C10, HM-727; Bifidobacterium adolescentis, Strain L2-32, HM-633; Clostridium innocuum, Strain 6_1_30, HM-173; Clostridium sp., Strain M62/1, HM-635; Bacteroides fragilis, Strain 3_1_12, HM-20; Bacteroides fragilis, Strain CL05T00C42, HM-711; Bacteroides fragilis, Strain CL07T00C01, HM-709; Bacteroides dorei, Strain CL02T00C15, HM-717; Bacteroides dorei, Strain 5_1_36/D4, HM-29; Bacteroides cellulosilytious, Strain CL02T12C19, HM-726; Bacteroides finegoldi, Strain CL09T03C10, HM-727; Clostridium symbiosum, Strain WAL-14673, HM-319; Clostridium citroniae, Strain WAL-17108, HM-315; Peptoclostridium difficile, Strain CD3, NR-43546; Peptoclostridium difficile, Strain CD178, NR-43504; Peptoclostridium difficile, Strain CD160, NR-43516; Clostridium difficile, Strain P7, NR-32887; Clostridium difficile, Strain P8, NR-32888; Clostridium difficile, Strain P11, NR-32890; Clostridium difficile, Strain P29, NR-32903; Clostridium difficile, Isolate-7, NR-13433.

## REFERENCES

1. Koizumi, S. (2007). Human heme oxygenase-1 deficiency: a lesson on serendipity in the discovery of the novel disease. Pediatr. Int. 49, 125–132.

2. Kenny, D.J., Plichta, D.R., Shungin, D., Koppel, N., Hall, A.B., Fu, B., Vasan, R.S., Shaw, S.Y., Vlamakis, H., Balskus, E.P., et al. (2020). Cholesterol Metabolism by Uncultured Human Gut Bacteria Influences Host Cholesterol Level. Cell Host Microbe 28, 245–257.e6.

3. Kang, D.-J., Ridlon, J.M., Moore, D.R., Barnes, S., and Hylemon, P.B. (2008). Clostridium scindens baiCD and baiH genes encode stereo-specific 7α/7β-hydroxy-3-oxo-Δ4-cholenoic acid oxidoreductases. Biochimica et Biophysica Acta (BBA) - Molecular and Cell Biology of Lipids 1781, 16–25.

4. Gérard, P. (2013). Metabolism of cholesterol and bile acids by the gut microbiota. Pathogens 3, 14–24.

5. Midtvedt, A.C., and Midtvedt, T. (1993). Conversion of cholesterol to coprostanol by the intestinal microflora during the first two years of human life. J. Pediatr. Gastroenterol. Nutr. 17, 161–168.

6. Midtvedt, T. (1974). Microbial bile acid transformation. Am. J. Clin. Nutr. 27, 1341–1347.

7. Ridlon, J.M., Harris, S.C., Bhowmik, S., Kang, D.-J., and Hylemon, P.B. (2016). Consequences of bile salt biotransformations by intestinal bacteria. Gut Microbes 7, 22–39.

8. Fahmy, K., Gray, C.H., and Nicholson, D.C. (1972). The reduction of bile pigments by faecal and intestinal bacteria. Biochim. Biophys. Acta 264, 85–97.

9. Ching, S., Ingram, D., Hahnel, R., Beilby, J., and Rossi, E. (2002). Serum levels of micronutrients, antioxidants and total antioxidant status predict risk of breast cancer in a case control study. J. Nutr. 132, 303–306.

10. Kapitulnik, J. (2004). Bilirubin: an endogenous product of heme degradation with both cytotoxic and cytoprotective properties. Mol. Pharmacol. 66, 773–779.

11. Poland, R.L., and Odell, G.B. (1971). Physiologic jaundice: the enterohepatic circulation of bilirubin. N. Engl. J. Med. 284, 1–6.

12. Brodersen, R., and Hermann, L.S. (1963). Intestinal reabsorption of unconjugated bilirubin. A possible contributing factor in neonatal jaundice. Lancet 1, 1242.

13. Vítek, L., Zelenka, J., Zadinová, M., and Malina, J. (2005). The impact of intestinal microflora on serum bilirubin levels. J. Hepatol. 42, 238–243.

14. Vítek, L., Kotal, P., Jirsa, M., Malina, J., Cerná, M., Chmelar, D., and Fevery, J. (2000). Intestinal colonization leading to fecal urobilinoid excretion may play a role in the pathogenesis of neonatal jaundice. J. Pediatr. Gastroenterol. Nutr. 30, 294–298.

15. Saxerholt, H., Carlstedt-Duke, B., Høverstad, T., Lingaas, E., Norin, K.E., Steinbakk, M., and Midtvedt, T. (1986). Influence of antibiotics on the faecal excretion of bile pigments in healthy subjects. Scand. J. Gastroenterol. 21, 991–996.

16. Lester, R., and Schmid, R. (1963). Intestinal Absorption of Bile Pigments. New England Journal of Medicine 269, 178–182. 10.1056/nejm196307252690402.

17. Levitt, D.G., and Levitt, M.D. (2014). Quantitative assessment of the multiple processes responsible for bilirubin homeostasis in health and disease. Clin. Exp. Gastroenterol. 7, 307–328.

18. Lester, R., and Schmid, R. (5 1965). Intestinal Absorption of Bile Pigments. III. The Enterohepatic Circulation of Urobilinogen in the Rat. J. Clin. Invest. 44, 722–730.

19. Saxerholt, H., Skar, V., and Midtvedt, T. (1990). HPLC separation and quantification of bilirubin and its glucuronide conjugates in faeces and intestinal contents of germ-free rats. Scand. J. Clin. Lab. Invest. 50, 487–495.

20. Gustafsson, B.E., and Lanke, L.S. (1960). Bilirubin and urobilins in germfree, ex-germfree, and conventional rats. J. Exp. Med. 112, 975–981.

21. Midtvedt, T., and Gustafsson, B.E. (1981). Microbial conversion of bilirubin to urobilins in vitro and in vivo. Acta Pathol. Microbiol. Scand. B 89, 57–60.

22. Hamoud, A.-R., Weaver, L., Stec, D.E., and Hinds, T.D. (2018). Bilirubin in the Liver–Gut Signaling Axis. Trends Endocrinol. Metab. 29, 140–150.

23. Leníček, M., Duricová, D., Hradsky, O., Dušátková, P., Jirásková, A., Lukáš, M., Nachtigal, P., and Vítek, L. (2014). The relationship between serum bilirubin and Crohn’s disease. Inflamm. Bowel Dis. 20, 481–487.

24. Schieffer, K.M., Bruffy, S.M., Rauscher, R., Koltun, W.A., Yochum, G.S., and Gallagher, C.J. (2017). Reduced total serum bilirubin levels are associated with ulcerative colitis. PLoS One 12, e0179267.

25. Vítek, L., Majer, F., Muchová, L., Zelenka, J., Jirásková, A., Branný, P., Malina, J., and Ubik, K. (2006). Identification of bilirubin reduction products formed by Clostridium perfringens isolated from human neonatal fecal flora. J. Chromatogr. B Analyt. Technol. Biomed. Life Sci. 833, 149–157.

26. Rekittke, I., Wiesner, J., Röhrich, R., Demmer, U., Warkentin, E., Xu, W., Troschke, K., Hintz, M., No, J.H., Duin, E.C., et al. (2008). Structure of (E)-4-hydroxy-3-methyl-but-2-enyl diphosphate reductase, the terminal enzyme of the non-mevalonate pathway. J. Am. Chem. Soc. 130, 17206–17207.

27. Hubbard, P.A., Liang, X., Schulz, H., and Kim, J.-J.P. (2003). The crystal structure and reaction mechanism of Escherichia coli 2,4-dienoyl-CoA reductase. J. Biol. Chem. 278, 37553–37560.

28. Dennery, P.A., Seidman, D.S., and Stevenson, D.K. (2001). Neonatal hyperbilirubinemia. N. Engl. J. Med. 344, 581–590.

29. Lloyd-Price, J., Arze, C., Ananthakrishnan, A.N., Schirmer, M., Avila-Pacheco, J., Poon, T.W., Andrews, E., Ajami, N.J., Bonham, K.S., Brislawn, C.J., et al. (2019). Multi-omics of the gut microbial ecosystem in inflammatory bowel diseases. Nature 569, 655–662.

30. Brink, M.A., Slors, J.F., Keulemans, Y.C., Mok, K.S., De Waart, D.R., Carey, M.C., Groen, A.K., and Tytgat, G.N. (1999). Enterohepatic cycling of bilirubin: a putative mechanism for pigment gallstone formation in ileal Crohn’s disease. Gastroenterology 116, 1420–1427.

31. Fevery, J. (1999). Pigment gallstones in Crohn’s disease. Gastroenterology 116, 1492–1494.

32. Lapidus, A., Akerlund, J.-E., and Einarsson, C. (2006). Gallbladder bile composition in patients with Crohn ‘s disease. World J. Gastroenterol. 12, 70–74.

33. Garrod, A.E. (1897). Note on the Origin of the Yellow Pigment of Urine. J. Physiol. 21, 190–191.

34. Weller, S.D.V. (1951). Bile pigments in the stools of infants. Arch. Dis. Child. 26, 86–88.

35. Maclagan, N.F. (1946). Faecal urobilinogen; clinical evaluation of a simplified method of estimation. Br. J. Exp. Pathol. 27, 190–200.

36. Keppler, D. (2014). The roles of MRP2, MRP3, OATP1B1, and OATP1B3 in conjugated hyperbilirubinemia. Drug Metab. Dispos. 42, 561–565.

37. Čvorović, J., and Passamonti, S. (2017). Membrane Transporters for Bilirubin and Its Conjugates: A Systematic Review. Front. Pharmacol. 8, 887.

38. Kamisako, T., Leier, I., Cui, Y., König, J., Buchholz, U., Hummel-Eisenbeiss, J., and Keppler, D. (1999). Transport of monoglucuronosyl and bisglucuronosyl bilirubin by recombinant human and rat multidrug resistance protein 2. Hepatology 30, 485–490.

39. Chen, S., and Tukey, R.H. (2018). Humanized UGT1 Mice, Regulation of UGT1A1, and the Role of the Intestinal Tract in Neonatal Hyperbilirubinemia and Breast Milk-Induced Jaundice. Drug Metab. Dispos. 46, 1745–1755.

40. Human Metabolome Database: Showing metabocard for Urobilinogen (HMDB0004158) https://hmdb.ca/metabolites/HMDB0004158.

41. Human Metabolome Database: Showing metabocard for Stercobilinogen (HMDB0004157) https://hmdb.ca/metabolites/HMDB0004157.

42. Naumann, H.N. (1947). Schlesinger’s test for urobilin in the presence of riboflavin and other fluorescent compounds. J. Lab. Clin. Med. 32, 1503–1507.

43. Elman, R., and McMaster, P.D. (1925). STUDIES ON UROBILIN PHYSIOLOGY AND PATHOLOGY: I. THE QUANTITATIVE DETERMINATION OF UROBILIN. J. Exp. Med. 41, 503–512.

44. Kotal, P., and Fevery, J. (1991). Quantitation of urobilinogen in feces, urine, bile and serum by direct spectrophotometry of zinc complex. Clin. Chim. Acta 202, 1–9.

45. Nordin, A., and Wagner, M. Application of 2D fluorescence spectroscopy on faecal pigments in water. https://stud.epsilon.slu.se/10251/1/daub_b_170622.pdf.

46. Miyabara, Y., Tabata, M., Suzuki, J., and Suzuki, S. (1992). Separation and sensitive determination of i-urobilin and 1-stercobilin by high-performance liquid chromatography with fluorimetric detection. J. Chromatogr. 574, 261–265.

47. Chao, A., and Goulding, C.W. (2019). A Single Mutation in the Mycobacterium tuberculosis Heme-Degrading Protein, MhuD, Results in Different Products. Biochemistry 58, 489–492.

48. Matsui, T., Nambu, S., Goulding, C.W., Takahashi, S., Fujii, H., and Ikeda-Saito, M. (2016). Unique coupling of mono-and dioxygenase chemistries in a single active site promotes heme degradation. Proceedings of the National Academy of Sciences 113, 3779–3784. 10.1073/pnas.1523333113.

49. Prjibelski, A., Antipov, D., Meleshko, D., Lapidus, A., and Korobeynikov, A. (2020). Using SPAdes De Novo Assembler. Curr. Protoc. Bioinformatics 70, e102.

50. Seemann, T. (2014). Prokka: rapid prokaryotic genome annotation. Bioinformatics 30, 2068–2069.

51. Kuznetsov, D., Tegenfeldt, F., Manni, M., Seppey, M., Berkeley, M., Kriventseva, E.V., and Zdobnov, E.M. (2023). OrthoDB v11: annotation of orthologs in the widest sampling of organismal diversity. Nucleic Acids Res. 51, D445–D451.

52. Dalkiran, A., Rifaioglu, A.S., Martin, M.J., Cetin-Atalay, R., Atalay, V., and Doğan, T. (2018). ECPred: a tool for the prediction of the enzymatic functions of protein sequences based on the EC nomenclature. BMC Bioinformatics 19, 334.

53. Jumper, J., Evans, R., Pritzel, A., Green, T., Figurnov, M., Ronneberger, O., Tunyasuvunakool, K., Bates, R., Žídek, A., Potapenko, A., et al. (2021). Highly accurate protein structure prediction with AlphaFold. Nature 596, 583–589.

54. Le Guilloux, V., Schmidtke, P., and Tuffery, P. (2009). Fpocket: an open source platform for ligand pocket detection. BMC Bioinformatics 10, 168.

55. Eberhardt, J., Santos-Martins, D., Tillack, A., and Forli, S. AutoDock Vina 1.2.0: New Docking Methods, Expanded Force Field, and Python Bindings. 10.26434/chemrxiv.14774223.

56. Trott, O., and Olson, A.J. (2010). AutoDock Vina: improving the speed and accuracy of docking with a new scoring function, efficient optimization, and multithreading. J. Comput. Chem. 31, 455–461.

57. Zhang, Y., and Skolnick, J. (2005). TM-align: a protein structure alignment algorithm based on the TM-score. Nucleic Acids Res. 33, 2302–2309.

58. Fu, L., Niu, B., Zhu, Z., Wu, S., and Li, W. (2012). CD-HIT: accelerated for clustering the next-generation sequencing data. Bioinformatics 28, 3150–3152.

59. Cantalapiedra, C.P., Hernández-Plaza, A., Letunic, I., Bork, P., and Huerta-Cepas, J. (2021). eggNOG-mapper v2: Functional Annotation, Orthology Assignments, and Domain Prediction at the Metagenomic Scale. Mol. Biol. Evol. 38, 5825–5829.

60. Parks, D.H., Chuvochina, M., Rinke, C., Mussig, A.J., Chaumeil, P.-A., and Hugenholtz, P. (2022). GTDB: an ongoing census of bacterial and archaeal diversity through a phylogenetically consistent, rank normalized and complete genome-based taxonomy. Nucleic Acids Res. 50, D785–D794.

61. Buchfink, B., Reuter, K., and Drost, H.-G. (2021). Sensitive protein alignments at tree-of-life scale using DIAMOND. Nat. Methods 18, 366–368.

62. Sievers, F., Wilm, A., Dineen, D., Gibson, T.J., Karplus, K., Li, W., Lopez, R., McWilliam, H., Remmert, M., Söding, J., et al. (2011). Fast, scalable generation of high-quality protein multiple sequence alignments using Clustal Omega. Mol. Syst. Biol. 7, 539.

63. Minh, B.Q., Schmidt, H.A., Chernomor, O., Schrempf, D., Woodhams, M.D., von Haeseler, A., and Lanfear, R. (2020). IQ-TREE 2: New Models and Efficient Methods for Phylogenetic Inference in the Genomic Era. Mol. Biol. Evol. 37, 1530–1534.

64. Foley, G., Mora, A., Ross, C.M., Bottoms, S., Sützl, L., Lamprecht, M.L., Zaugg, J., Essebier, A., Balderson, B., Newell, R., et al. (2019). Engineering indel and substitution variants of diverse and ancient enzymes using Graphical Representation of Ancestral Sequence Predictions (GRASP). bioRxiv. 10.1101/2019.12.30.891457.

65. Jones, P., Binns, D., Chang, H.-Y., Fraser, M., Li, W., McAnulla, C., McWilliam, H., Maslen, J., Mitchell, A., Nuka, G., et al. (2014). InterProScan 5: genome-scale protein function classification. Bioinformatics 30, 1236–1240.

66. Letunic, I., and Bork, P. (2021). Interactive Tree Of Life (iTOL) v5: an online tool for phylogenetic tree display and annotation. Nucleic Acids Res. 49, W293–W296.

67. Danecek, P., Bonfield, J.K., Liddle, J., Marshall, J., Ohan, V., Pollard, M.O., Whitwham, A., Keane, T., McCarthy, S.A., Davies, R.M., et al. (2021). Twelve years of SAMtools and BCFtools. Gigascience 10. 10.1093/gigascience/giab008.

68. Langmead, B., and Salzberg, S.L. (2012). Fast gapped-read alignment with Bowtie 2. Nat. Methods 9, 357–359.

