## Supplemental Information for "Discovery of the gut microbial enzyme responsible for bilirubin reduction to urobilinogen"

#### Figure S1

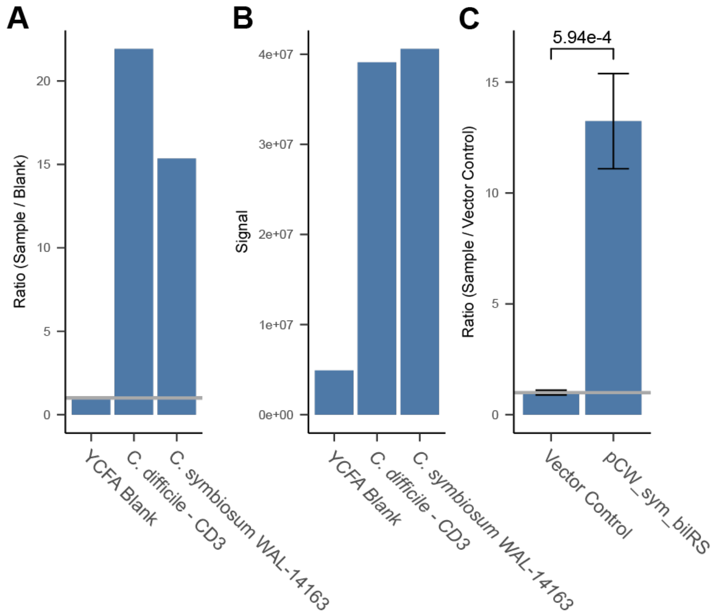


**Figure S1:** Confirmation of reduction of bilirubin and mesobilirubin to urobilin. A) Fluorescence assay comparing bilirubin reduction of *C. difficile* and *C. symbiosum* to the YCFA media blank (two-sample t-test). Bars show the ratio of the samples fluorescence to the corresponding experiments media blank. The gray line marks a ratio of 1, equivalent to the vector control. B) Metabolomics confirmation of urobilinogen production (two-sample t-test). C) Fluorescence assay comparing mesobilirubin reduction of ectopically expressed *C. symbiosum bilRS* in *E. coli* to the vector control (two-sample t-test). Bars show the ratio of the samples fluorescence to the corresponding experiments media blank. The gray line marks a ratio of 1, equivalent to the vector control.

Figure S2

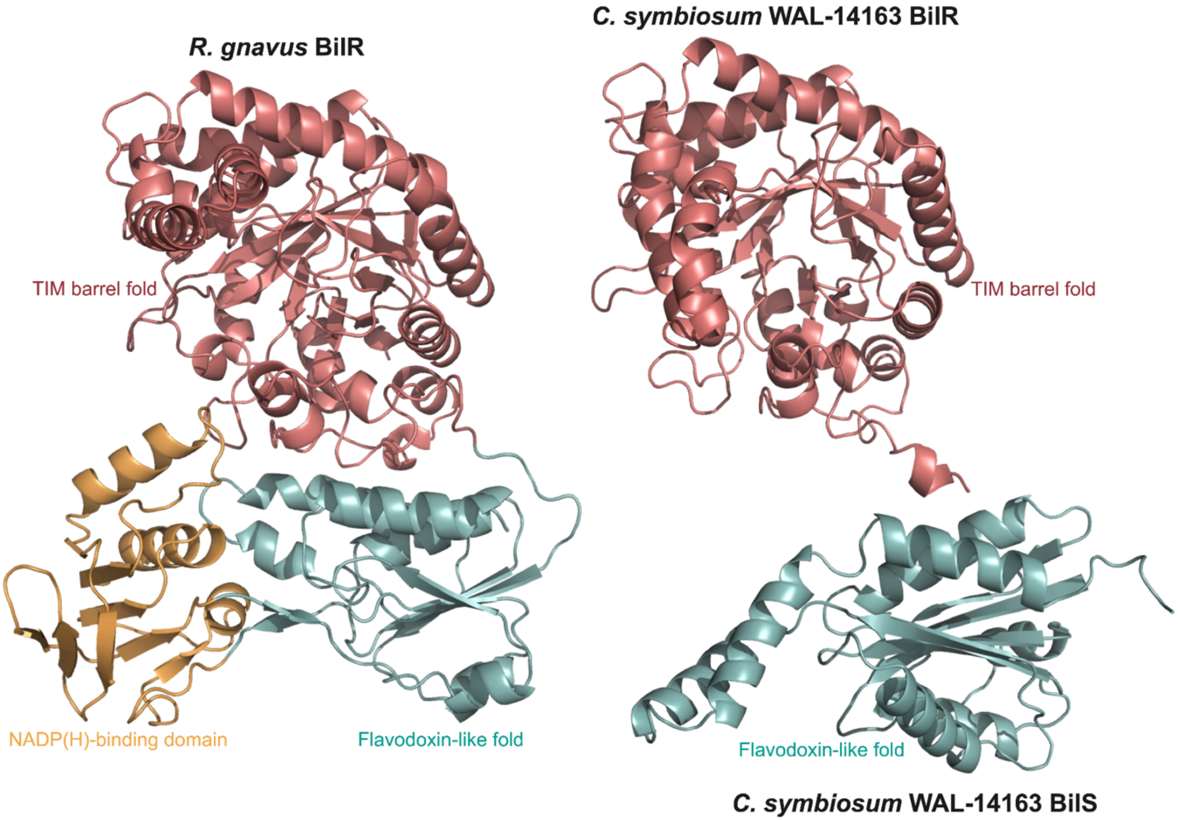


**Figure S2:** Predicted BilR and BilS structures. The *C. symbiosum* BilR and BilS were aligned to the *R. gnavus* BilR structure and are shown in the same orientation to show the similar fold structure. The structures are colored based on the predicted protein domains.

### Figure S3

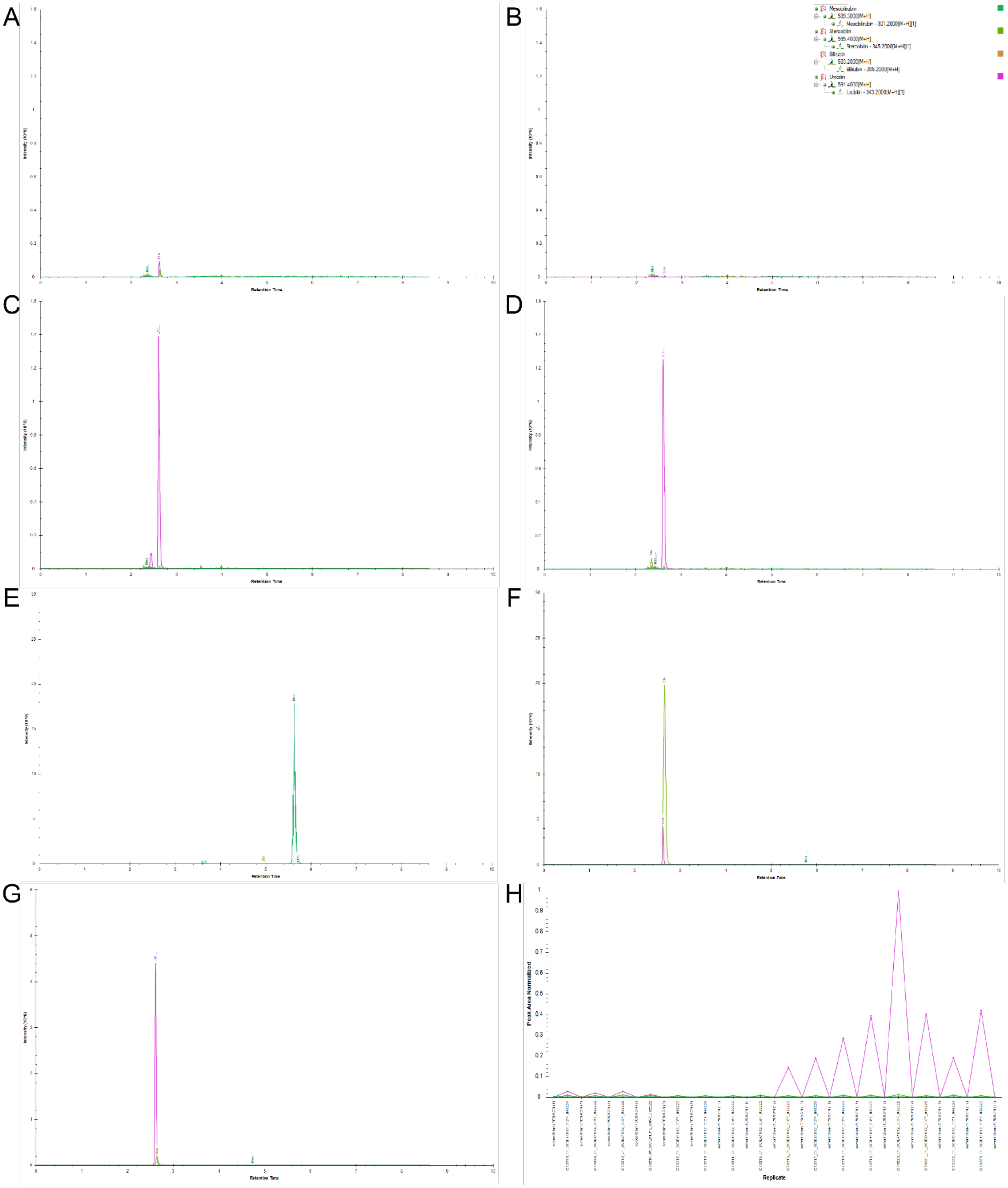
A
B
C
D
E
F
G
H
Mesobilirubin
589.3000[M+H]
Mesobilirubin - 301.2000[M+H][1]
Stercobilin
595.4000[M+H]
Stercobilin - 345.2000[M+H][1]
Bilirubin
583.2000[M+H]
Bilirubin - 285.2000[M+H]
Urobilin
591.4000[M+H]
Urobilin - 343.2000[M+H][1]
Retention Time
Peak Area Normalized
Replicate

**Figure S3:** (A-G) Extraction ion chromatograms for Mesobilirubin (m/z 589.3 →m/z 301.2) Urobilin (m/z 591.4→ m/z 343.2) and Stercobilin (m/z 595.4→m/z 345.2) in a BHI blank Sample (A) pCW-lic vector control sample (B), *C. symbiosum* sample (C), *C. difficile* sample (D), Bilirubin-Mesobilirubin standard (E), Stercobilin standard (F), and Urobilin standard (G). (H) The calculated peak area of each of the compounds in the samples.

Figure S4

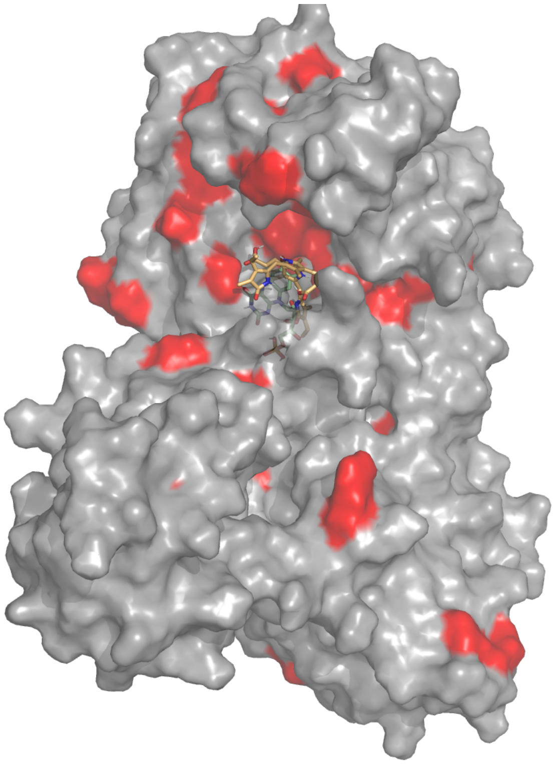

**Figure S4:** Changing and conserved positions in BilR structure. Predicted *R. gnavus* BilR structure with docked bilirubin (orange) and FMN (green) structures shown as stick models. Residues on the BilR structure are colored red if the position was conserved in more than 75% of the BilR sequences and if the probability of the most likely amino acid changed by more than 50% in the BilR ancestral node based on the ancestral sequence reconstruction.

Figure S5

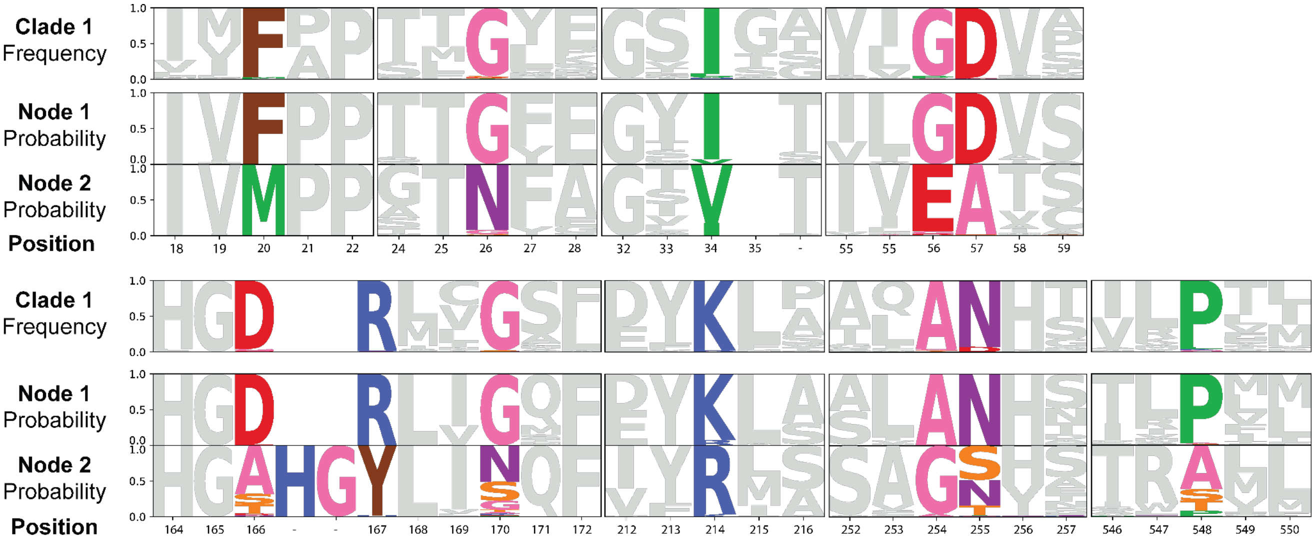


**Figure S5:** Residues of interest in BilR sequence. Sequence logos showing the conservation within the BilR clade sequences (top) and probability of amino acids in the BilR ancestral node (middle) and ancestor of other sequences (bottom). Positions are colored if they showed greater than 60% change in the probability of the most common amino acid between the two ancestral sequences and were conserved in greater than 90% of BilR sequences.

Figure S6

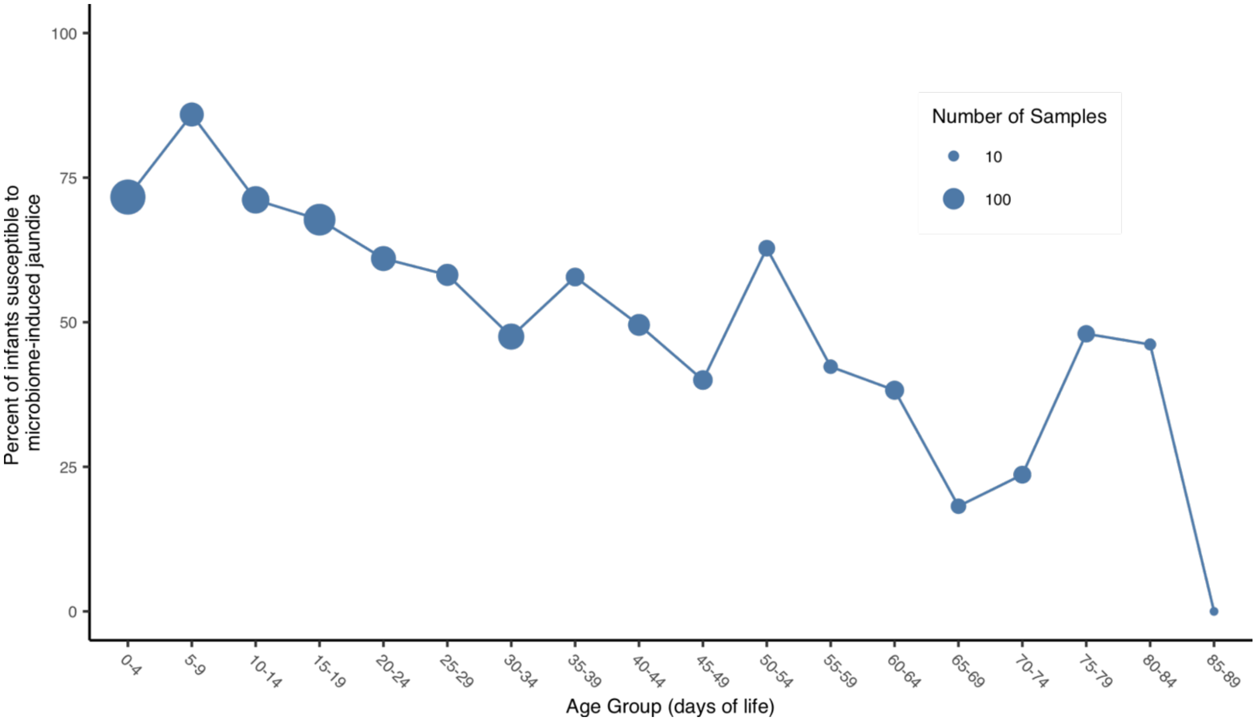


**Figure S6:** Frequent absence of bilR during the first three months of life. Plot showing the percent of samples within each five day age bin with no bilirubin reductase detected. Points are sized based on the number of samples included in each bin.
